# Triggering visually-guided behavior by holographic activation of pattern completion neurons in cortical ensembles

**DOI:** 10.1101/394999

**Authors:** Luis Carrillo-Reid, Shuting Han, Weijian Yang, Alejandro Akrouh, Rafael Yuste

## Abstract

Neuronal ensembles are building blocks of cortical activity yet it is unclear if they have any causal role in behavior. Here we tested if the precise activation of neuronal ensembles with two-photon holographic optogenetics in mouse primary visual cortex alters behavioral performance in a visual task. Disruption of behaviorally relevant cortical ensembles by activation of non-selective neurons decreased behavioral performance whereas optogenetic targeting of as few as two neurons with pattern completion capability from behaviorally relevant ensembles improved task performance by reliably recalling the whole ensemble. Moreover, in some cases, activation of two pattern completion neurons, in the absence of visual stimulus, triggered correct behavioral responses. Our results demonstrate a causal role of neuronal ensembles in a visually guided behavior and suggest that ensembles could represent perceptual states.

## Introduction

Cortical circuits generate synchronous activity states, also known as neuronal ensembles (or assemblies), that may constitute emergent functional units, as building blocks of memories, percepts, movements, or mental states (Abeles, 1991; Buzsaki, 2010; Churchland et al., 2012; Hopfield, 1982; Villette et al., 2015; Yuste, 2015). In mouse visual cortex, visual stimuli activate groups of neurons with coordinated activity defining neuronal ensembles (Carrillo-Reid et al., 2015b; Cossart et al., 2003a; Miller et al., 2014). These ensembles are also present in spontaneous activity, indicating that they can be stored and replayed by cortical circuits (Carrillo-Reid et al., 2016; MacLean et al., 2005; Miller et al., 2014). Moreover, using two-photon optogenetics, artificial ensembles can be stably imprinted in awake animals and later recalled by stimulating individual neurons, demonstrating that cortical circuits have pattern completion capability (Carrillo-Reid et al., 2016). However, the functional role of recalled cortical ensembles in behavior, if any, still remains unclear.

To explore this, we combined calcium imaging of neuronal populations (Yuste and Katz, 1991), two-photon microscopy (Denk et al., 1990; Yuste and Denk, 1995) and population analysis (Carrillo-Reid et al., 2017; Carrillo-Reid et al., 2015a) to identify neuronal ensembles in primary visual cortex from awake mice performing a visually guided Go/No-Go behavioral task. Then, using two-photon holographic optogenetics (Nikolenko et al., 2008; Packer et al., 2015; Rickgauer et al., 2014; Yang et al., 2018), we activated specific groups of neurons overlapped with visual stimuli to disrupt or recall cortical ensembles, while measuring the effect on behavioral performance. The use of a simple Go/No-Go task allowed us to precisely study changes in behavioral performance evoked by photostimulation at different contrast levels of visual stimuli. We take advantage of the existence of pattern completion neurons to manipulate neuronal ensembles optogenetically. We show that optogenetic activation of random group of cells during normal contrast visual stimuli disrupted cortical ensembles and deteriorated behavior whereas specific activation of neurons with pattern completion capability reliably recalled behaviorally relevant ensembles and improved task performance with low contrast visual stimuli. Moreover, optogenetic targeting of behaviorally relevant pattern completion neurons could even triggered behavior in the absence of visual stimulus.

## Results

### Head-fixed mice reliably perform visual Go/No-Go task

We carried out simultaneous two-photon imaging (GCaMP6s) and two-photon holographic optogenetics (C1V1) of targeted neurons in layer 2/3 of primary visual cortex (Packer et al., 2015; Rickgauer et al., 2014; Yang et al., 2018) through a reinforced thinned-skull window (Drew et al., 2010) in awake head-fixed mice that have been trained in a Go/No-Go visually guided task consisting of orthogonal drifting-gratings (Figs. 1A and 1B). Mice underwent a regime of habituation to the treadmill and water restriction for 2 days until they reached 85% of their original weight. After this habituation period, mice went through 3 days of continuous reinforcement where water reward was delivered following the Go signal (at 100% visual contrast). After this continuous reinforcement period, and to avoid sudden changes in pupil diameter due to high contrast visual stimuli, we reduced the contrast level to 50%. During this training protocol mice gradually learned to lick correctly when Go and No-Go visual stimuli were randomly presented. After 7 days of performing the visually-guided behavioral task (at 50% contrast) mice reached a performance level above 75% that plateau for at least 8 days. We considered expert mice those with a behavioral performance above 75% from day 10 on (Fig. 1C; Performance = hits/(hits+miss) – false choices/(false choices+correct rejects). Improvement in behavioral performance (Fig. 1D; day 1: 31±5%; expert: 97±1%; P<0.005**) due to increased hits (day 1: 83±7%; expert: 99±1%; P<0.005**) and reduced false choices (day 1: 52±8%; expert: 3±1%; P<0.005**) was accompanied by a faster licking onset (Fig. 1E; day 1: 1711±84s; expert: 988±146s; P<0.005**). Low contrast levels of visual stimuli (10% - 40%) generated a reduction of behavioral performance (Fig. 1F; normal contrast: 82±4%; low contrast: 54±4%; P<0.005**). These experiments demonstrated that head-fixed mice can perform consistently a visually guided Go/No-Go task.

**Fig. 1.**
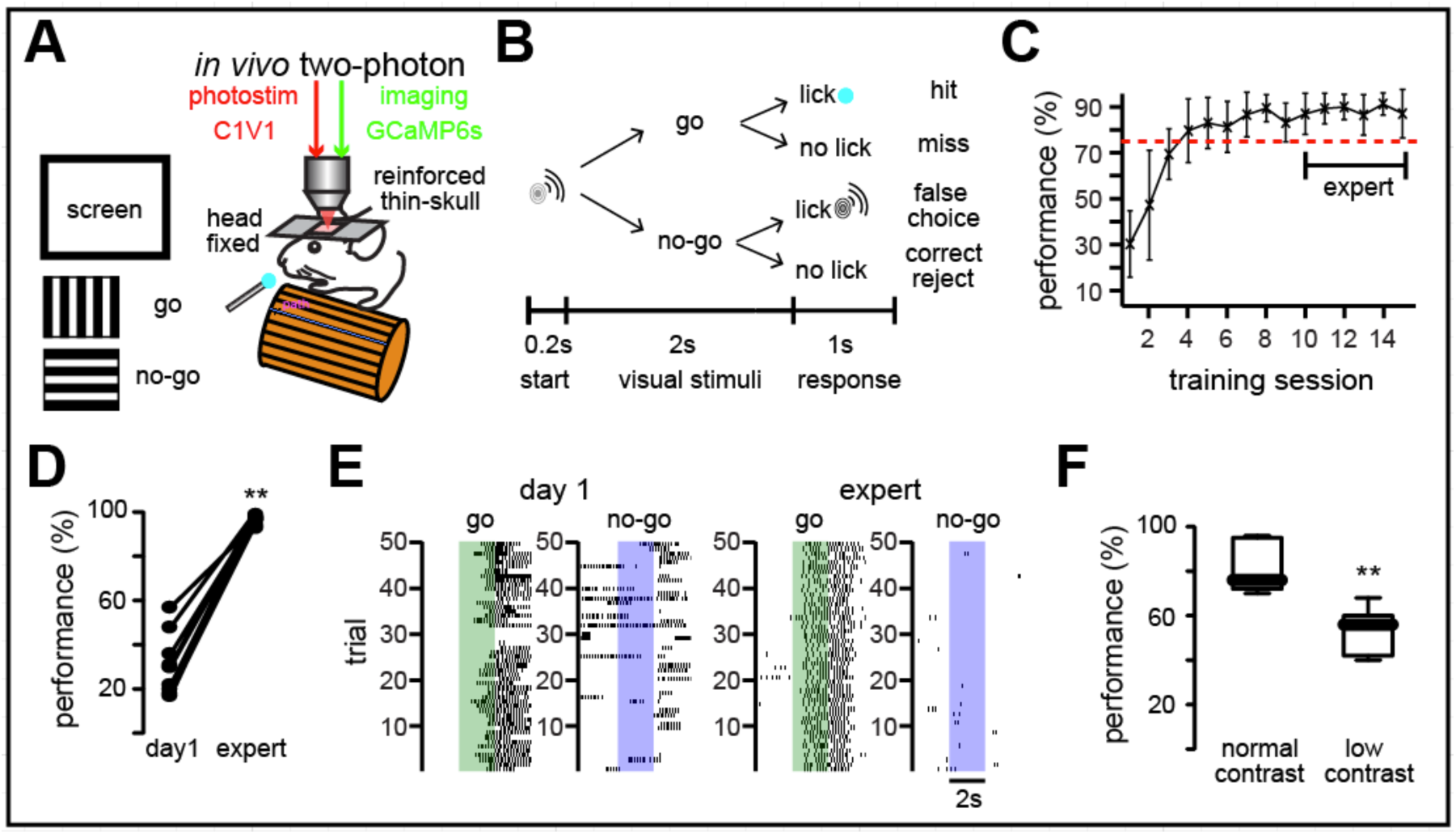
Visually-guided Go/No-Go task. (**A**) Experimental design: simultaneous two-photon calcium imaging and two-photon optogenetic manipulation of targeted neurons in visually guided Go/No-Go task. (**B**) Performance assessment. (**C**) Improvement in performance as a function of training session. (**D**) Performance increased significantly in expert mice (P<0.005**; n = 9 mice; Wilcoxon matched-pairs signed rank test). (**E**) Enhancement of behavioral performance was reflected as shorter licking delays. Colored bars represent visual stimuli (Go: green; No-Go: blue; expert: day 10). Dark markers correspond to lick. (**F**) Worsening in performance by low contrast visual stimuli in expert animals (P<0.005**; n = 7 mice. Data presented as whisker box plots displaying median and interquartile ranges analyzed using Mann-Whitney test).

### Identification of neuronal ensembles and pattern completion neurons

We followed previous studies, which have shown activation of neuronal ensembles by visual stimuli consisting of different orientations of drifting-gratings in layer 2/3 of primary visual cortex (Carrillo-Reid et al., 2015b; Miller et al., 2014). To identify neuronal ensembles we first turned the changes in fluorescence into a digital raster plot of activity. Mathematically, neuronal ensembles can be understood as multidimensional population vectors where each vector indicates the joint activation of a neuronal population at a different point in time (Fig. 2A). The dimensionality of the ensembles corresponds to the total number of imaged neurons. Because the same neuron can respond to multiple orientations, we searched for clusters in this multidimensional space to identify neuronal ensembles that responded to the same visual stimuli. Population vectors indeed formed clusters in a multidimensional space (Fig. 2B). We first visualized clusters defining neuronal ensembles using principal component analysis (PCA) as a commonly used multidimensional reduction technique (Carrillo-Reid et al., 2016). To compare these clusters quantitatively we quantified the normalized inner product between population vectors and used factorization of similarity matrices of the normalized inner product of all possible vector pairs. Since similarity matrices are symmetric, we then used singular value decomposition (SVD) to rigorously identify potential neuronal ensembles (Carrillo-Reid et al., 2015a; Carrillo-Reid et al., 2015b). After the identification of the ensembles we used a conditional random field (CRF) model (Fig. 2C) to find the neurons that are most representative for each ensemble, based on their predictability and the node strength of functional connections between neurons (Fig. 2D) (Carrillo-Reid et al., 2017). Such neurons could be optically targeted for two-photon optogenetic stimulation (Fig. 2E) using a spatial light modulator (SLM) (Nikolenko et al., 2008). Indeed, neurons with high functional connectivity have pattern completion capabilities (Carrillo-Reid et al., 2016), so the identification of these pattern completion “critical” neurons can enable the targeted optical manipulation of neuronal ensembles (Carrillo-Reid et al., 2017).

**Fig. 2.**
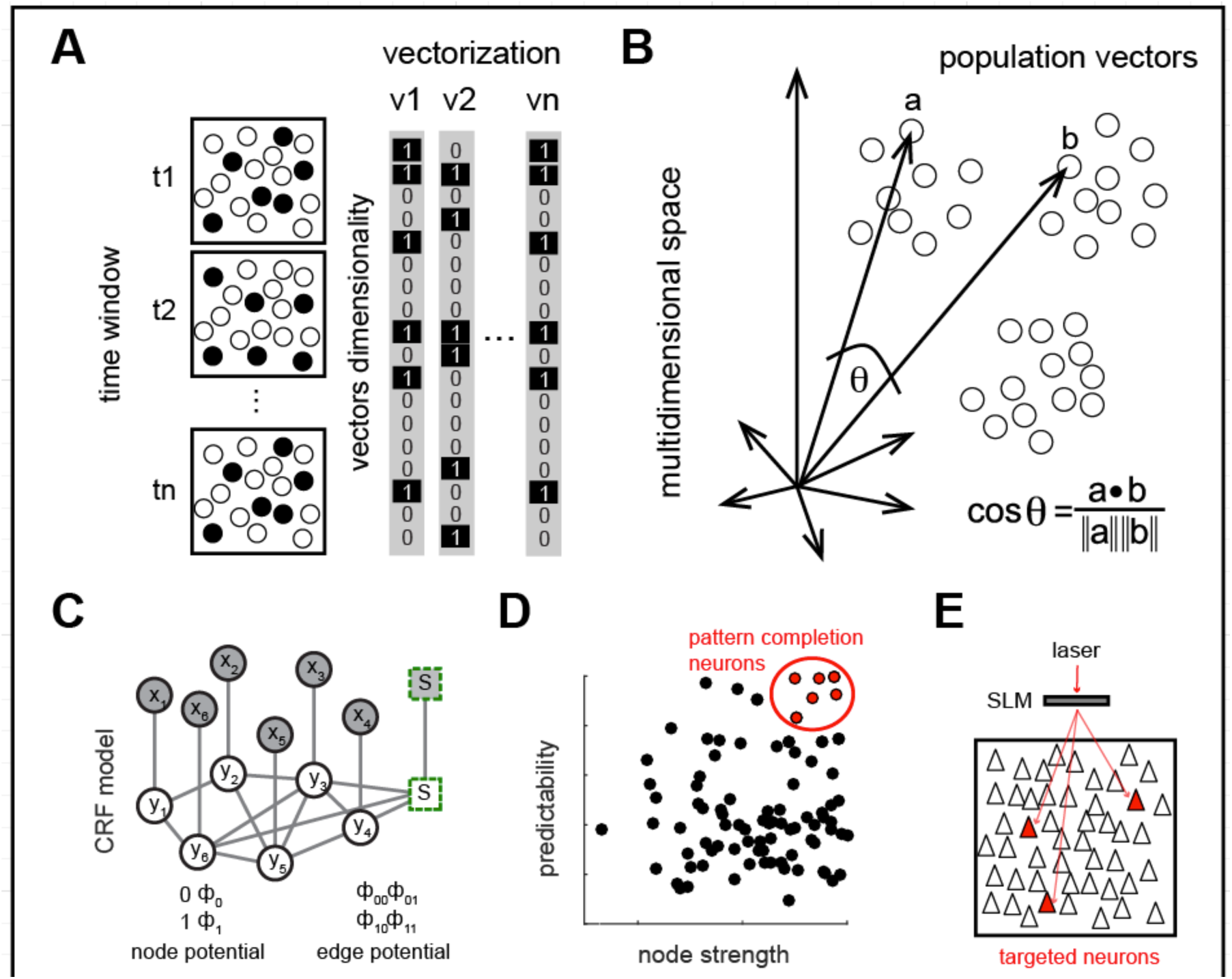
Identification of neuronal ensembles and neurons with pattern completion capabilities. (**A**) Schematic representation of neuronal activity at different time points and their representation as population vectors (**B**) Cartoon of population vectors in a multidimensional space. Each dot represents one population vector and clusters of population vectors define a neuronal ensemble. The normalized inner product compares population vectors by the cosine of the angle between any pair of vectors in a multidimensional space. (**C**) Graphical representation of Conditional Random Field (CRF) models. Circles represent neurons. Visual stimulus is represented by an added node (square). Shaded nodes (x) represent observed data. White nodes (y) represent neurons from the graphical model. Edges indicate the mutual probabilistic dependencies between neurons. Node potentials indicate if a neuron is active or inactive. Edge potentials represent states of adjacent neurons. (**D**) Identification of most representative neurons from cortical ensembles, related to a given visual stimuli, defined by predictability values computed as the AUC from the ROC curve and node strengths (top right neurons). Red are the neurons with pattern completion capability. (**E**) Neurons with pattern completion capability that co-express GCaMP6s and C1V1 can be simultaneously photostimulated using a SLM.

### Reliably activation of Go-signal neuronal ensembles after training

We then searched for Go and No-Go neuronal ensembles by performing PCA of population vector activity (Carrillo-Reid et al., 2016) in expert mice during behavior. Indeed, PCA showed the presence of a single Go-signal neuronal ensemble, which differed from separate clusters of population vectors representing different neuronal ensembles related to No-Go stimuli (Fig. 3A). SVD factorization (Carrillo-Reid et al., 2015b; Carrillo-Reid et al., 2016) confirmed the engagement of a neuronal ensemble that was reliably recalled during the Go signal, whereas No-Go visual stimuli recruited different set of population vectors visualized in the similarity map as different blocks of activity (Fig. 3B). The temporal course of ensemble activation computed by SVD factorization confirmed that neurons belonging to the Go-signal neuronal ensemble reliably responded to Go stimuli, whereas variable groups responded to No-Go signal (Fig. 3C) The spatial analysis of the activated neurons revealed that Go and No-Go neuronal ensembles constituted non-overlapping neuronal subgroups, and that population vectors evoked by No-Go visual stimuli fluctuate at different time points (Fig. 3D). To quantify the similarity between population vectors evoked by Go and No-Go signals we computed the normalized inner product between all the population vectors that belong to the Go ensemble and compared them against all the population vectors evoked by No-Go visual stimuli demonstrating that population vectors from the Go ensemble differed from No-Go signals (Fig. 3E; similarity index Go: 0.38±0.0055; similarity index Go vs No-Go: 0.046±0.0018; P<0.0001). To quantify neuronal ensemble reliability we computed the percentage of times that a given visual stimuli activated a group of neurons above chance levels from the total number of Go or No-Go presentations. This demonstrated that, in expert mice, the Go ensemble was reliably activated when the Go signal was presented. On the contrary, No-Go visual stimuli poorly recalled its associated neuronal ensembles (Fig. 3F; reliability go: 88±4; reliability no-go: 38±4; P<0.005**). In addition, a significantly lower amount of coactive neurons were recalled by No-Go visual stimuli, as compared to the Go signal (Fig. 3G; coactive neurons Go: 7.6±0.2; coactive neurons No-Go: 2.9±0.3; P<0.0001****). This suggests that, in expert mice, Go ensembles become more reliable and the responsiveness of cortical microcircuits to No-Go visual stimuli is somewhat suppressed. However, the number of representative neurons defining Go and No-Go ensembles, albeit lower, was not significantly different (Fig. 3H; ensemble neurons Go: 14.1±1.5; ensemble neurons No-Go:12.3±1.1; P>0.05 n.s.), as previously shown for neuronal ensembles representing different orientations (Carrillo-Reid et al., 2015b; Carrillo-Reid et al., 2016; Miller et al., 2014). These experiments demonstrate that, in trained mice, Go neuronal ensembles are activated by visual stimulation more reliably than other neuronal ensembles.

**Fig. 3.**
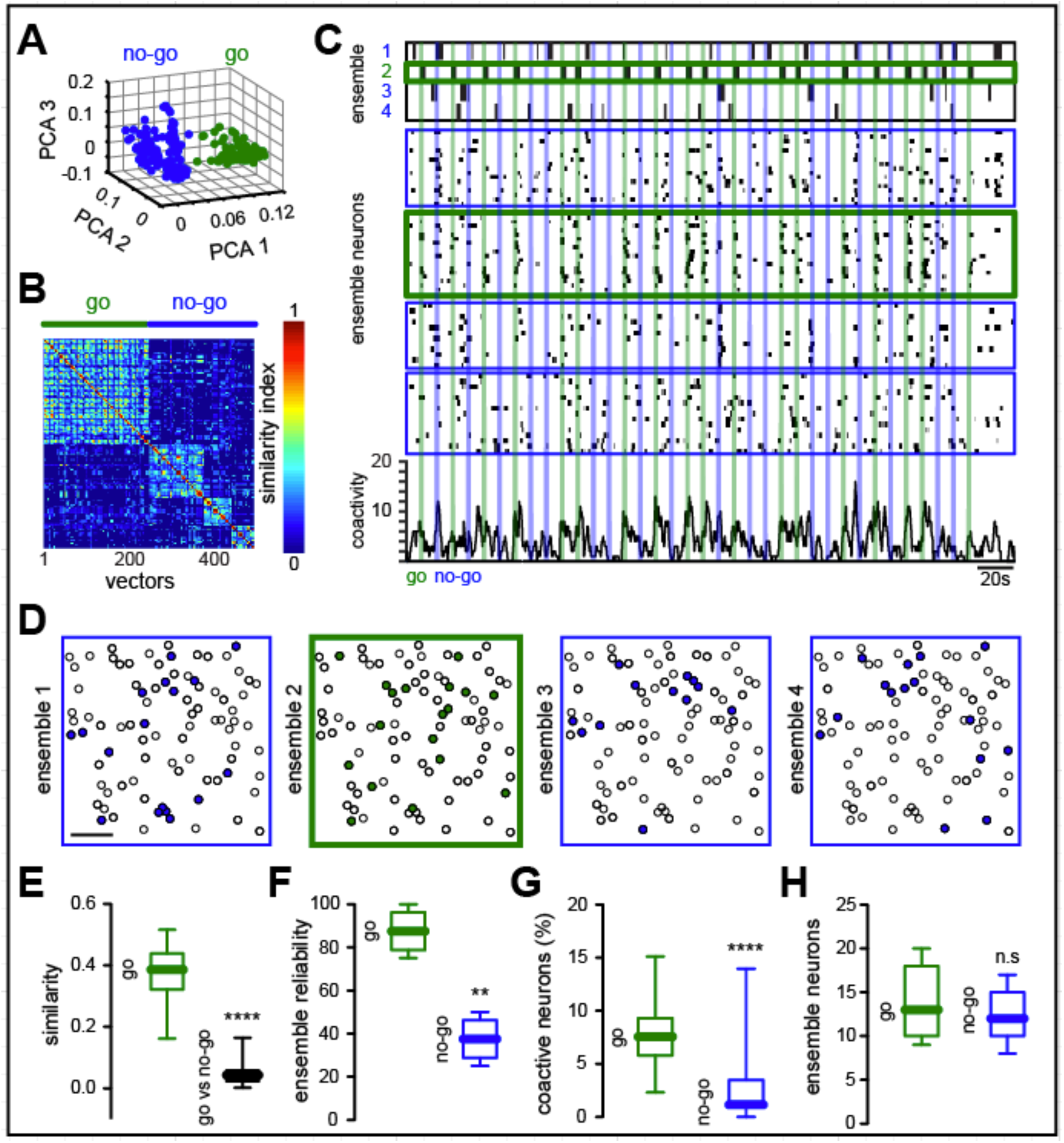
Reliable activation of neuronal ensemble by Go stimulus. (**A**) Different ensembles activated in Go and No-Go tasks in trained mice. PCA of population vectors evoked by visual stimuli show that coactive groups of neurons responding to the “Go” signal (green) define a cluster of vectors that differs from those activated by the “No-Go” signal (No-Go: blue). Each dot represents a population vector. (**B**) Sorted similarity map representing lack of overlap between population vectors from Go and No-Go ensembles. (**C**) Top: Time course of ensembles identified with SVD (Green: Go; Blue: No-Go). Middle: Raster plot of neurons belonging to these four neuronal ensembles (same order as in top). Note variability in individual responses. Bottom: Histogram of activity from all recorded neurons. Note that some No-Go trials have reduced network activity. (**D**) Spatial maps of same data showing that different subsets of neurons belong to the four cortical ensembles. Scale bar 50 µm. (**E**) Cosine similarity between population vectors related to Go and No-Go stimulus (P<0.0001). (**F**) Reliability of Go ensembles is higher than that of No-Go ensembles (P<0.005). (**G**) Number of coactive neurons is reduced during No-Go stimuli (P<0.0001). (**H**) Number of neurons from Go and No-Go ensembles is similar (P>0.05). Data presented as whisker box plots displaying median and interquartile ranges analyzed using Mann-Whitney test.

### Two-photon imaging of holographic activation of targeted neurons

After finding that a specific group of neurons are reliably recalled by behaviorally relevant Go visual stimuli we wondered if the activation of a handful of targeted neurons could alter behavioral performance. To test this, we used two-photon holographic patterns created by a spatial light modulator (SLM) to optogenetically target selective sets of neurons simultaneously (Packer et al., 2015; Rickgauer et al., 2014; Yang et al., 2018). To perform simultaneous two-photon imaging and two-photon optogenetics in targeted groups of neurons we used a holographic microscope with two independent two-photon lasers, one to image GCaMP6s (940 nm) and another to activate the red shifted opsin C1V1 (1040 nm) (Yang et al., 2018). Co-expression of GCaMP6s and C1V1 only occurred in ∼50% of the neurons (Fig. 4A). To test if the joint photostimulation of several neurons was spatially restricted we targeted different combinations of pyramidal neurons in layer 2/3 of primary visual cortex and monitored the calcium transients of adjacent neurons (Figs. 4B and 4C). Targeted neurons co-expressing GCaMP6s and C1V1 showed clear changes in fluorescence evoked by photostimulation, compared to non-targeted neurons (Fig. 4D; fluorescence targeted: 34±3%; fluorescence non-targeted: 4±0.1%; P<0.0001****) demonstrating that our approach can be used to selectively activate particular neuronal populations.

**Fig. 4.**
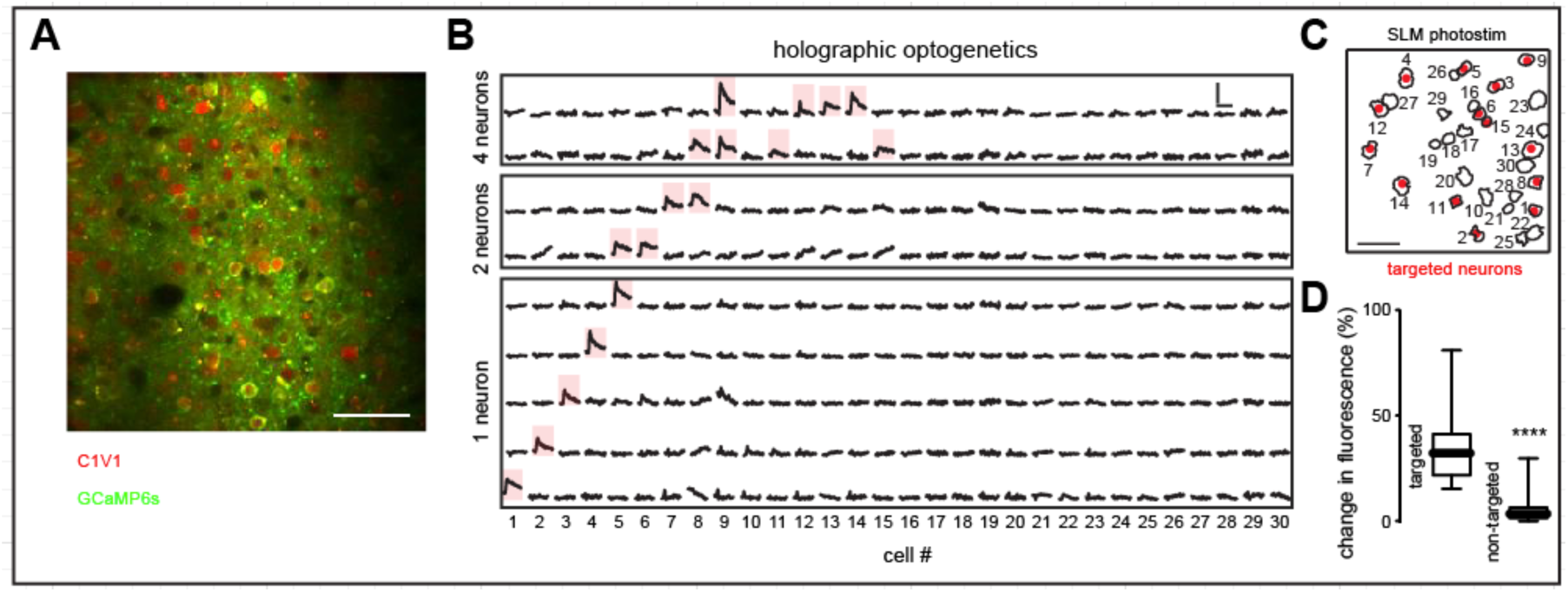
Simultaneous two-photon optogenetic photostimulation of cortical neurons. (**A**) Representative field of view depicting neurons co-expressing C1V1-mCherry (red) and GCaMP6s (green). Note that co-expression is sparse. (**B**) Calcium transients from neurons co-expressing C1V1 and GCaMP6s when different subsets of targeted cells were photostimulated. Red shadows show reliably responsive neurons when one or multiple cells were targeted using a Spatial Light Modulator (SLM). Scale bars: 10 sec and 50% change in fluorescence. (**C**) Spatial map of neurons co-expressing C1V1 and GCaMP6s. Targeted cells shown in **A** are highlighted in red. Scale bar 50 µm. (**D**) Changes in fluorescence evoked in targeted neurons were significantly different than non-targeted neurons (n=30 neurons; P<0.0001; Mann Whitney test). Data presented as whisker box plots displaying median and interquartile ranges.

### Holographic activation of non-GO neurons disrupts ensemble identity and behavioral performance

To test the link between neuronal ensembles and behavior we proceeded in three steps. First, we activated, during the Go signal, a nonspecific group of neurons that did not belong to the Go ensemble (Fig. 5A; “Disrupt” condition = Go stimulus + SLM stimulation). This manipulation degraded the identity of the Go ensemble, creating a mixed response, visualized as population vectors that clearly differed from visually evoked neuronal ensembles (Fig. 5B). Accordingly, the similarity map of population vectors evoked by visual stimuli and population vectors evoked by the Disrupt condition revealed two different populations (Fig. 5C). Population analysis demonstrated that population vectors evoked by Go-signal were significantly different from those population vectors during the Disrupt condition (Figs. 5D; similarity index go vs. disrupt: 0.033±0.0036; similarity index Disrupt: 0.43±0.0066; P<0.0001****), confirming that the Go ensemble were disrupted by the optogenetic stimulation by the targeted activation of a random group of neurons. SVD vector factorization showed different neuronal ensembles defining Go, No-Go and Disrupt ensembles. Specifically, the activity from Go and No-Go ensemble neurons was significantly reduced when the Disrupt neurons were activated together with visual stimuli (Fig. 5E). Disrupt ensembles (whose neurons were chosen randomly) were mostly composed by neurons not belonging to Go or No-Go ensembles (Fig 5F and 5G; not belonging neurons: 9.50±1.1; Go neurons: 2.5±0.4; No-Go neurons: 2.2±0.3; P<0.005**). The Disrupt protocol lead to a reduced reliability of Go ensemble activation (Fig. 5H; Go ensemble reliability in Disrupt conditions: 4.3±1%; P<0.005**) and reduced the cross-correlation between neurons belonging to Go ensembles (Fig. 5I; cross-correlation go: 0.27±0.0149; cross-correlation disrupt: 0.02±0.0053; P<0.005**). Together with these changes in the Go ensemble, the Disrupt condition also led to significant decreases in task performance (Fig. 5J; performance Go: 81.5±4%; performance Disrupt: 66.8±6%; P<0.05*) and increased onsets for licking (Fig. 5K; lick onset go: 1.2±0.1938s; lick onset disrupt: 1.6±0.2558s; P<0.05*). These experiments demonstrate that the targeted activation of a handful group of neurons can influence behavioral performance. We conclude that the Go ensemble is *necessary* for the correct execution of the visually guided task, since the disruption of the Go ensemble by the optogenetic activation of non-specific neurons was accompanied by a degradation of the behavior.

**Fig. 5.**
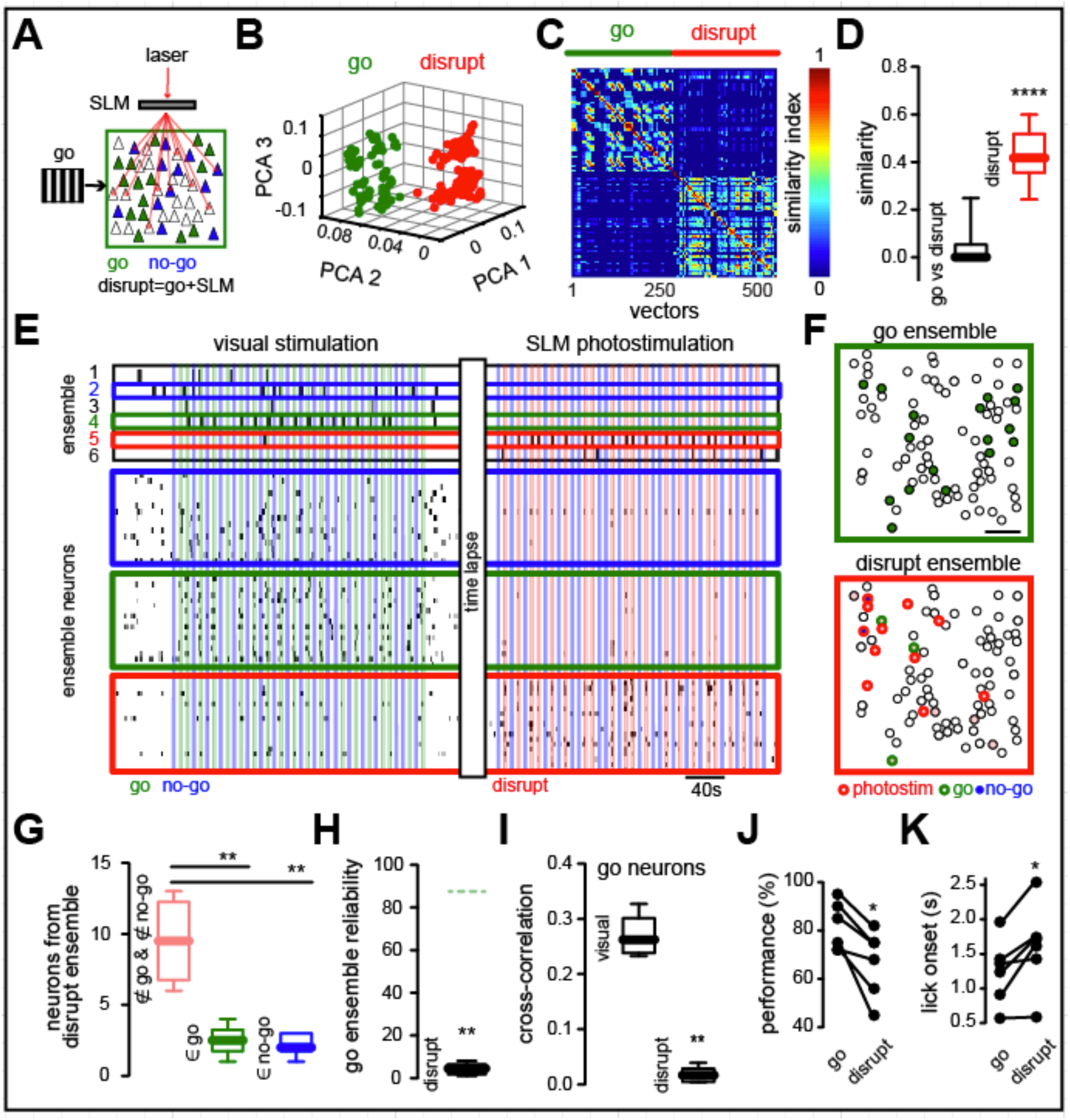
Unspecific neuronal activation degrades ensemble identity and visual performance. (**A**) Experimental protocol. Green neurons represent Go ensemble; blue represent No-Go ensemble. During the Disrupt condition, unspecific sets of neurons (red; including neurons from No-Go ensemble) are simultaneously photostimulated during Go stimulus presentation. (**B**) Disruption of Go ensemble identity by stimulation of Disrupt neurons. PCA of population vectors evoked by “Go” stimulus alone and with simultaneous photoactivation of disrupt neurons, which generates a different cortical response (red). Each dot represents a population vector. (**C**) Similarity maps of multidimensional population vectors showing that photostimulation of Disrupt neurons during Go visual stimuli breaks down Go ensemble identity. (**D**) Go and Disrupt ensembles are significantly different (P<0.0001). (**E**) Top: Neuronal ensemble analysis shows the creation of an artificial neuronal ensemble (red) by targeted activation of Disrupt neurons. Bottom: Raster plots of neurons belonging to Go, No-Go and Disrupt ensembles. Note reduction of neuronal responses to the “Go” and “No-Go” neurons evoked by the simultaneous activation of Disrupt neurons. (**F**) Spatial map of neurons from Go (green) and Disrupt ensembles (red). Optogenetic targeting included neurons belonging to No-Go ensemble. Scale bar 50 µm. (**G**) Disrupt ensemble is composed mainly of optogenetically targeted neurons (P<0.005). (**H**) Reliability of Go ensemble during disruption is significantly decreased during activation of disrupt neuron (P<0.005). Green dotted line represents Go ensemble reliability in control conditions. (**I**) Cross-correlation of Go ensemble neurons is significantly reduced by Disrupt protocol (P<0.005). (**J**) Behavioral performance is significantly decreased during Disrupt protocol (P<0.05). Data presented as whisker box plots displaying median and interquartile ranges analyzed using Mann-Whitney test. (**K**) Licking onset is significantly increased by Disrupt protocol (P<0.05). n = 6 mice; Wilcoxon matched-pairs signed rank test.

### Activation of Go ensembles by holographic optogenetics of pattern completion neurons improves behavioral performance

In a second step, we investigated whether the targeted recalling of the Go ensemble could improve behavioral performance, by holographic optogenetic activation of Go ensemble neurons during behavior (Fig. 6A). To do so, we first decreased the contrast of visual stimuli in trained mice in order to reduce task performance (Glickfeld et al., 2013), thereby increasing our sensitivity to detect behavioral changes in trained animals. Under low contrast visual stimulation conditions (10-40% contrast), the behavioral performance of trained animals significantly decreased (Fig. 1F). Given our past finding that stimulation of one or a few pattern completion neurons can recall an entire ensemble (Carrillo-Reid et al., 2016), we chose to selectively target several of them for photostimulation, using our CRF graph theory method to identify them first computationally from their responses to the Go stimulus (Fig. 2D; (Carrillo-Reid et al., 2017)). In order to perform these set of experiments, neurons with pattern completion capability must also co-express GCaMP6s and C1V1, so only a few of the animals used in the present study satisfied the criteria (6/122 mice) and very few pattern completion neurons were available for stimulation.

**Fig. 6.**
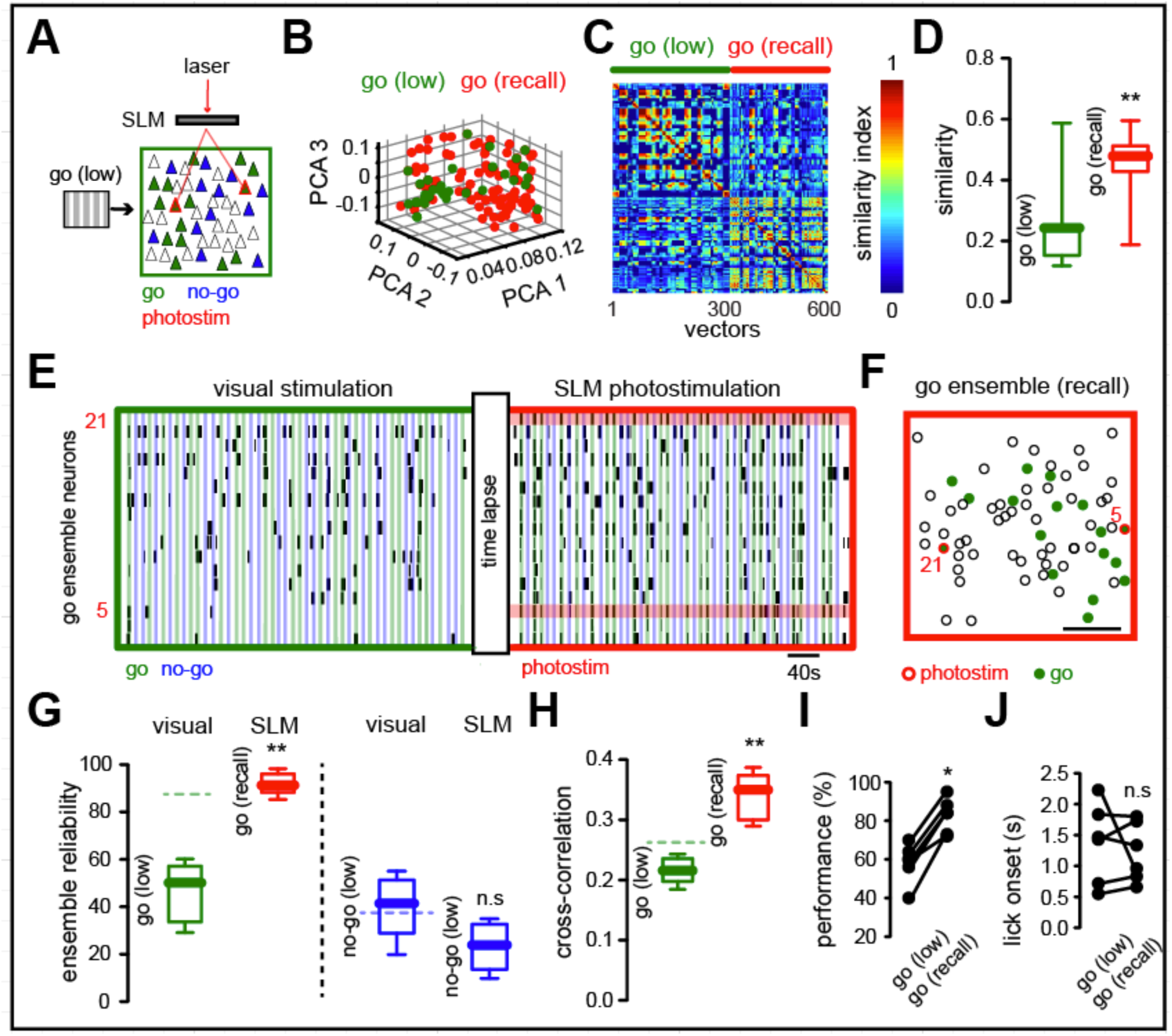
Activation of pattern completion neurons reliably improves Go ensemble and enhances task performance. (**A**) Experimental protocol. Go stimulus neurons (green) in low contrast conditions and simultaneously targeted pattern completion neurons (red) that belonged to the Go ensemble (Recall condition). (**B**) PCA of population vectors evoked by the low contrast Go stimulus (green) and simultaneous low contrast Go visual stimulation and activation of pattern completion neurons (red). Each dot represents a population vector. (**C**) Similarity maps of population vectors representing the Go ensemble in low contrast condition alone and with simultaneous holographic photostimulation. (**D**) Recall condition increases Go ensemble reliability. (**E**) Raster plot from neurons belonging to the Go ensemble shows change in overall activity evoked by the simultaneous activation of two neurons with pattern completion capability (neurons 5 and 21; red bars). Note that the reliability of individual neuronal responses is increased. (**F**) Spatial map of layer 2/3 neurons highlighting neurons belonging to the Go ensemble (green). SLM photostimulated neurons in red. Scale bar 50 µm. (**G**) Reliability of Go and No-Go ensembles during visual stimulation and SLM photostimulation of pattern completion neurons belonging to the Go ensemble. Left: the reliability of recalled Go ensemble is significantly increased from Go ensemble in low contrast conditions (P<0.005). Right: the reliability of No-Go ensemble remains unaltered (P>0.05). Green and blue dotted lines represent the mean values from Go and No-Go ensemble reliability in control conditions respectively. (**H**) The cross-correlation of neurons belonging to the Go ensemble is significantly increased by SLM targeting of neurons with pattern completion capability (P<0.005). Data presented as whisker box plots displaying median and interquartile ranges analyzed using Mann-Whitney test. (**I**) Behavioral response to low contrast Go-Signal is significantly enhanced by the targeted activation of pattern completion neurons (P<0.05). (**J**) The mean value of the licking onset was not significantly reduced under Recall conditions (P>0.05). n = 6 mice; Wilcoxon matched-pairs signed rank test.

Holographic activation of two or more pattern completion neurons during low contrast visual stimuli generated evoked population vectors that overlapped with those originally evoked by Go-signals, confirming their pattern completion capabilities (Fig 6B). Similarity maps depicting the angles between population vectors belonging to the Go ensemble and Go ensembles activated by holographic photostimulation of pattern completion neurons in the presence of Go signals in low contrast indicated that both ensembles were indistinguishable (Fig. 6C). Consistent with this, the similarity between population vectors was significantly increased by photostimulation (Fig. 6D; similarity index go (low): 0.26±0.0578; similarity index go (recall): 0.46±0.0074; P<0.005**), demonstrating that optogenetic targeting of several pattern completion neurons could be used reliably to activate (or “Recall”) a neuronal ensemble that represents a behaviorally relevant stimulus. The raster plot of neurons during the Recall condition showed that Go ensemble neurons were more reliably activated during optogenetic targeting of pattern completion neurons (Fig. 6E). Recall Go ensembles had a widespread spatial distribution and pattern completion neurons were not spatially clustered (Fig. 6F). As indicated by the similarity map (Fig. 6C) and raster plots (Fig. 6E), the reliability of Go ensembles in low contrast stimuli was significantly lower than that of Recall Go ensembles (Fig. 6G; left; reliability Go (low contrast): 46.7±5%; reliability Go (Recall): 91.5±2%; P<0.005**). No-Go ensemble reliability remained unaltered by holographic stimulation of neurons with pattern completion capability (Fig. 6G; right; reliability No-Go (low contrast; no optogenetics): 40±5%; reliability No-Go (low contrast with optogenetics): 23±4%; P>0.05 n.s). The increase in Go ensemble reliability during the Recall condition was reflected as enhanced cross-correlation of Go ensemble neurons (Fig 6H; cross-correlation go (low contrast): 0.22±0.0089; cross-correlation go (Recall): 0.34±0.0155; P<0.005**). Consistent with this, the targeted optogenetic manipulation of pattern completion neurons significantly improved behavioral performance (Fig. 6I; performance Go (low): 58.3±4%; performance go (Recall): 82.6± 3.6%; P<0.05*). Even though there was a shortening of the licking onset, this was not significant (Fig. 6J; lick onset Go (low): 1.37±0.2623s; lick onset go (Recall): 1.22±0.1949s; P>0.05 n.s). These results demonstrate a correlation between the increase in reliability of the Go ensemble by stimulation of pattern completion neurons and the enhancement of behavioral performance of a visually guided behavior.

### Stimulation of at least two pattern completion neurons is necessary to enhance behavioral response

We wondered whether the behavioral performance could also be enhanced by the photoactivation of a single pattern completion neuron, since artificially imprinted ensembles, formed by the repetitive activation of a random group of neurons, can be recalled by single cell stimulation (Carrillo-Reid et al., 2016). To test this, we used two-photon optogenetics to activate one pattern completion neuron during low contrast visual stimuli (Fig. 7A), finding that this evoked population vectors that overlapped those evoked by activation of multiple pattern completion neurons (Fig. 7B). Moreover, photostimulation of individual pattern completion neurons also evoked population vectors that were similar to population vectors evoked by Go signals with low contrast (Fig. 7C; similarity single: 0.34±0.0226; P>0.05 n.s). However, the total number of recalled neurons was significantly enhanced by photostimulation of multiple neurons (Fig. 7D; recalled neurons from single neuron activation: 4±0.52; recalled neurons multiple neuron activation: 6.68±0.44; P<0.005**). Indeed, the activation of several pattern completion neurons was more effective in recalling behaviorally relevant neuronal ensembles (Fig. 7E). Recalled ensembles after single neuron stimulation were also distributed across the field of view (Fig. 7F) and the cross-correlation of the recalled neurons was not significantly different from those of visual stimulation alone (Fig. 7G; cross-correlation single: 0.24±0.0114; P>0.05 n.s). Ensemble reliability also remained the same between ensembles evoked by low contrast visual stimuli and single cell activation (Fig. 7H; reliability single: 50±5%; P>0.05 n.s). Consistent with this, the behavioral performance and the hit percentage were not enhanced by single neuron activation compared to low contrast visual stimuli, whereas activation of multiple pattern completion neurons still enhanced behavioral performance (Fig. 7I; performance Go (low contrast): 58.3±4%; performance single: 66.3±3.6%; performance SLM: 82.6±4%; P(low vs single)>0.05 n.s; P(single vs SLM)<0.005**) by an increase in the number of hits (Fig. 7J; hits go (low): 76±3.4%; hits single: 77.7±3.4%; hits SLM: 89.5±3.3%; P(low vs single)>0.05 n.s; P(single vs SLM)<0.05*). Thus, in these experiments, the targeted activation of a single pattern completion neuron was not able to reliably recall a Go-ensemble or enhance behavioral performance of a visually-guided task. We concluded that, under our experimental conditions, the holographic activation of at least two neurons with pattern completion capability is required to generate significant effects in behavioral performance.

**Fig. 7.**
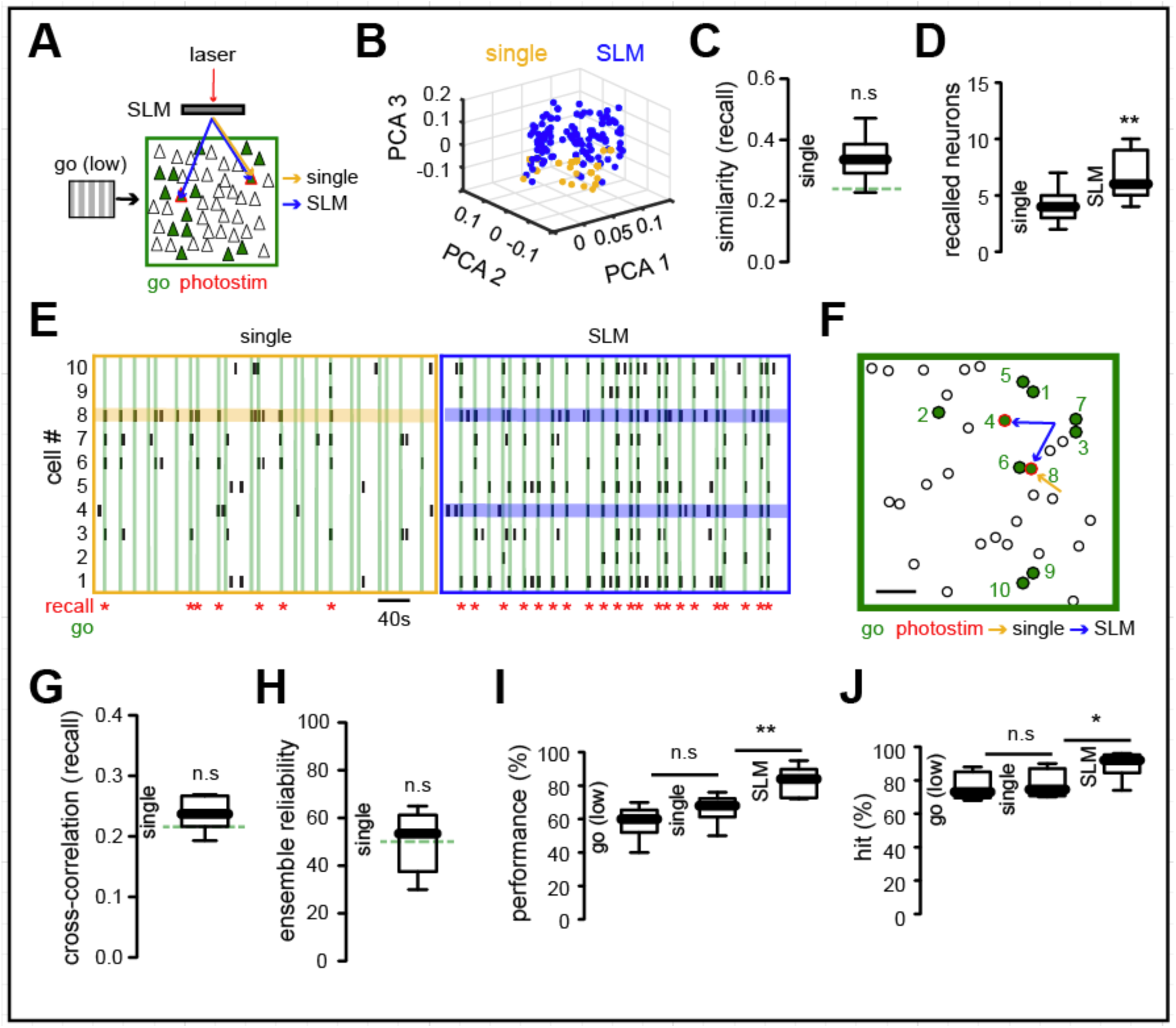
Stimulation of more than one pattern completion neuron is necessary to improve behavioral performance. **(A)** Schematic representation of experimental conditions. Optogenetic stimulation of individual neurons with pattern completion capability was performed simultaneously with the presentation of low contrast Go stimuli. One (orange) or, for comparison, multiple neurons (blue) were stimulated. **(B)** PCA of population vectors evoked by single (orange) or multiple pattern completion neurons (blue). Each dot represents a population vector. (**C**) Population vectors evoked by single cell photostimulation are not significantly different from population vectors evoked by Go stimuli (green). (**D**) Simultaneous photostimulation of multiple pattern completion neurons increased the number of recalled neurons (P<0.005**). (**E**) Raster plot of the most representative neurons belonging to Go ensemble. Horizontal lines highlight photostimulated neurons (orange: single cell stimulation; blue: simultaneous photostimulation of multiple neurons). Recalled ensembles highlighted in red. Note that activation of multiple pattern completion neurons reliably recalls the Go ensemble. (**F**) Spatial map of photostimulated and recalled ensemble. Scale bar 50 µm. (**G**) The cross-correlation of Go ensemble neurons (green) was not altered by stimulation of individual pattern completion neurons (P>0.05). (**H**) Go ensemble reliability (green) remained unaltered by stimulation of individual pattern completion neurons (P>0.05). Green lines represent mean values from Go ensembles (low). (**I**) Behavioral performance with single cell activation was not significantly different from low contrast visual stimuli alone (P>0.05) whereas activation of multiple neurons significantly increased behavioral performance (P< 0.005**). (**J**) Number of correct hits was not significantly different between single cell stimulation and visual stimuli alone (P>0.05) whereas it was significantly increased by multiple neuron stimulation (P<0.05*). n= 6 mice. Data presented as whisker box plots displaying median and interquartile ranges using Mann-Whitney test.

### Behavioral responses triggered by ensemble activation by pattern completion neurons in the absence of visual stimulus

To continue examining the behavioral role of ensembles, in a third step, we trigger the Go ensemble capability in the absence of any visual stimulation, by stimulating two pattern completion neurons from the Go ensemble together (Fig. 8A). In some instances, the dual stimulation led to the recalling of the Go ensemble and this was accompanied by a major increase in behavioral performance, compared to the trials when the Go ensemble was not recalled (Fig. 8B; performance no recall: 18.3±2.8%; performance recall: 70.8±3.5%; P<0.005**). This suggests that optogenetic activation of multiple pattern completion neurons substituted for the Go stimulus. Consistent with this, the licking onset evoked by the recalling of the Go ensemble in the absence of visual stimuli was not significantly different from the licking onset evoked by Go visual stimuli under normal conditions (50% contrast) (Fig. 8C; licking onset visual: 1.2±0.1938s; licking onset recall: 1.6±0.1608s; P>0.05 n.s). The cross-correlation of neurons belonging to the Go ensemble was also significantly higher during successful recalling epochs compared to non-recalling epochs (Fig. 8D; cross-correlation no recall: 0.12±0.0064; cross-correlation recall: 0.26±0.0144; P<0.005**), indicating that Go ensemble neurons were activated together during the recalling epochs, as can be directly seen in the raster plot from neurons belonging to the Go ensemble (Fig. 8E). Recalled Go ensembles in the absence of visual stimuli also had a widespread spatial distribution (Fig. 8F) and the number of recalled neurons without visual stimuli in each trial was similar to the number of neurons recalled by visual stimuli (Fig. 8G; recalled neurons SLM: 6.5±1.1; P>0.05 n.s). These experiments demonstrate that in trained mice, stimulation of pattern completion neurons successfully triggered the Go ensemble and the correct behavioral response.

**Fig. 8.**
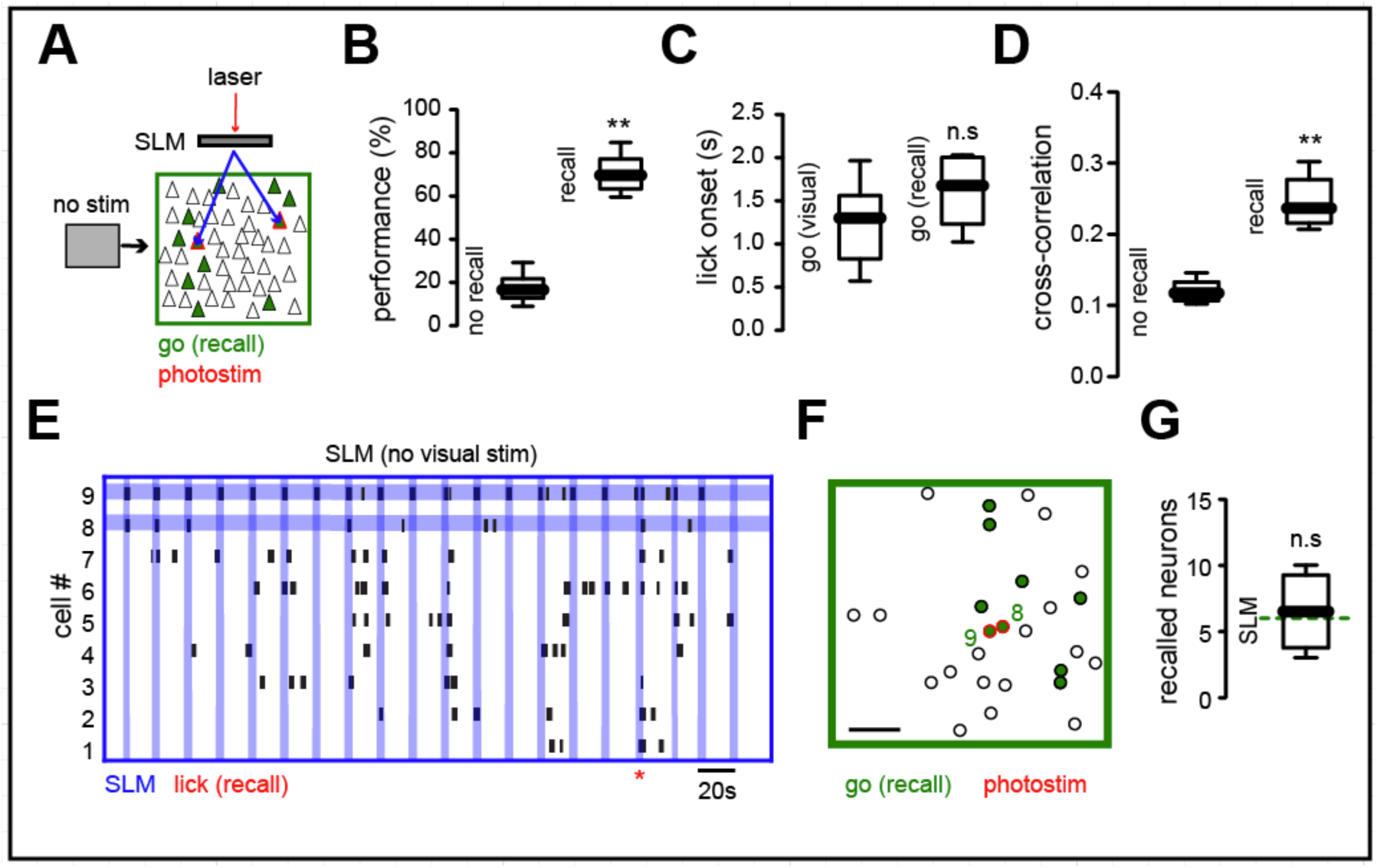
Behavior induced by recalling Go ensemble in absence of visual stimuli. (**A**) Experimental conditions. Simultaneous optogenetic stimulation of pattern completion neurons was performed in the absence of visual stimuli (animals viewed a gray screen). (**B**) Behavioral performance evoked by recalling the Go ensemble by optogenetic stimulation in the absence of visual stimuli was significantly higher than performance in non-recall trials (P<0.005**). (**C**) Licking onset from successfully driven optogenetic behavioral events was not significantly different from licking onset in visual evoked behavior (P>0.05). n= 6 mice. (**D**) Paired cross-correlation of Go ensemble neurons was enhanced during successful recall, compared with non recall trials (P<0.005**). (**E**) Raster plot of most representative neurons from Go ensemble during holographic stimulation of pattern completion neurons. Vertical blue lines indicate photostimulation. Horizontal lines highlight targeted neurons. Red marker shows successful recalling of Go ensemble and licking behavior. (**F**) Spatial map of E showing stimulated and recalled neurons during successful licking trial. (**G**) Number of recalled neurons after optogenetic stimulus was not significantly different from active neurons in low contrast conditions (line). Data presented as whisker box plots displaying median and interquartile ranges using Mann-Whitney test.

## Discussion

Here we report that the precise holographic activation of targeted neurons can selectively and bidirectionally modify behavioral performance, and even substitute for the visual stimulus altogether, demonstrating that neuronal ensembles in layer 2/3 of mouse primary visual cortex constitute functional cortical units. Alterations in ensemble identity generated by targeted two-photon optogenetics disrupted behavioral performance in predictable ways: while activation of neurons not related to the perceptual task degraded behavioral performance, the precise activation of a behaviorally meaningful neuronal ensemble enhanced the behavior elicited by low contrast visual stimuli or could even trigger the behavior in the absence of visual stimulation. These experiments demonstrated that cortical ensembles are necessary and sufficient for visually-guided behavior.

### Pattern completion in neocortical circuits

Pattern completion, defined as the ability to recall a complex pattern of information from a small part of it, is a cornerstone of human memory and many behaviors. In a neural circuit, pattern completion is thought to occur when an initial activity pattern is imprinted in a set of neurons via the strengthening of its connections (Seung and Yuste, 2010). After this stage, the activation of only one the neurons sets off the entire group. Initially proposed by Marr to explain associative recall in the hippocampus (Marr, 1971), the intrinsic ability of recurrently connected neural circuits to generate pattern completion has been highlighted by theorists, helping the system converge on attractors states (Hopfield, 1982; Hopfield and Tank, 1986). Completion of patterns of spikes was first described in hippocampus, using electrophysiological recordings (Mizumori et al., 1989), and it has been suggested that CA3, with its recurrent connectivity, may play a particularly important role in implementing it (Gold and Kesner, 2005). More recently, hippocampal pattern completion has been linked to visual discrimination in the cortex (Hindy et al., 2016), indicating that pattern completion may not be specific to the hippocampus but found widely throughout the forebrain. In agreement with this, we previously found that coactivation of a group of neurons could imprint them to fire together as an ensemble, and that, for days after this, the stimulation of a single neuron could trigger the activation of the entire ensemble, a clear and direct demonstration of pattern completion by a neural circuit (Carrillo-Reid et al., 2016). This discovery implies that cortical circuits can function by the activation of modules, composed of groups of neurons, and that these modules are controlled by a few selected cells who can trigger them. Because of this useful property, and regardless of the mechanisms and its functional significance, in this study we use pattern completion as a tool to activate neuronal ensembles. As we demonstrate (Figures 6, 7 and 8), activation of pattern completion neurons is an effective method to externally manipulate the activity of neuronal populations, and, due to this property, we speculate that pattern completion will be a key mechanism internally used by neural circuits to activate neuronal ensembles.

### Comparison with previous findings

Previous reports using electrical stimulation (Afraz et al., 2006; Bartlett and Doty, 1980; Brecht et al., 2004; DeAngelis et al., 1998; Doty, 1965; Gu et al., 2012; Romo et al., 1998; Salzman et al., 1990) or one-photon optogenetic stimulation (Huber et al., 2008) of neurons in different cortical areas have reported behavioral correlates of neuronal activation. Our results, based on the precise manipulation and observation of functional ensembles, indicate that such observed effects, including the reports of individual neurons triggering motor responses (Brecht et al., 2004), could be due to the recalling of specialized neuronal ensembles by pattern completion neurons (Carrillo-Reid et al., 2016). Indeed, the recalling of neuronal ensembles related to the Go signal only produced a significant enhancement of behavioral performance when at least two neurons with pattern completion capability were simultaneously activated (Fig. 7). Moreover, the fact that we can bidirectionally enhance or deteriorate behavioral performance as a function of the targeted neurons indicates that animal responses after electrical microstimulation (Bartlett and Doty, 1980; Salzman et al., 1990) may critically depend on the accurate recalling of functional cortical circuits.

### Behavioral relevance of ensembles

One possible interpretation of our results would be that targeted optogenetics are just modulating the sensory stimulus at the cortical level. But perhaps the clearer demonstration of the functional importance of the ensemble is the fact that their activation triggered the behavioral responses in the absence of a visual stimulus (Fig. 8). This fascinating result suggests that the sensory stimulus can be substituted in trained animals by the activation of a neuronal ensemble, as opposed to the activation of individual neurons (Romo et al., 1998), perhaps thought a cascade of activation of downstream targets through pattern completion. Thus, the perception of a specific visual stimulus, as demonstrated by the correct behavior, can be internally driven and be independent of the sensory input. In this scenario, ensembles could be viewed as dynamical attractors that implement perceptual or memory states (Hopfield, 1982), rather than mere sensory states. Indeed, the ability to generate states of activity that are independent of the sensory realm, and which can be used to symbolize or mentally manipulate the world, has been long suspected to underlie the design logic of many areas of the central nervous system (Hebb, 1949; Hopfield, 1982; Lorente de No, 1938).

Finally, the demonstration that the precise manipulation of cortical ensemble identity can selectively alter behavioral performance opens the possibility to study the physiological role of targeted functional circuits with single cell resolution in awake behaving animals in other brain areas and learned tasks. The development of holographic microscopy (Yang et al., 2018) and animal models co-expressing opsins and genetically encoded calcium indicators, could help discern the exact role of emergent states of activity, such as neuronal ensembles, as modular building blocks of neural circuits during functional or pathological behavioral states.

## Acknowledgments

Laboratory members for help and virus injections. Stanford Neuroscience Gene Vector and Virus Core for AAVdj virus. We thank Clay Lacefield for advice in the behavioral setup. Supported by the National Eye Institute (DP1EY024503, R01EY011787), National Institute of Mental Health (R01MH101218; R01MH100561) and Defense Advanced Research Projects Agency (SIMPLEX N66001-15-C-4032). This material is based upon work supported by, or in part by, the U. S. Army Research Laboratory and the U. S. Army Research Office (Contract W911NF-12-1-0594, MURI). The authors declare no competing financial interests. L.C.-R. and R.Y. conceptualized this work. L.C.R., S.H., W.Y. and A.A. analyzed the data. L.C.-R., W.Y. and A.A. performed the experiments. L.C.-R. wrote the original draft. L.C.-R. and R.Y. reviewed and edited the paper. R.Y. assembled and directed the team and secured funding. All the data are archived at the NeuroTechnology Center at Columbia University.

## Methods

### Animals

All experimental procedures were carried out in accordance with the US National Institutes of Health and Columbia University Institutional Animal Care and Use Committee. Experiments were performed on C57BL/6 male mice that were 60-90 days of age before headplate implantation. Animals were housed on a 12h light-dark cycle with food and water *ad libitum*.

### Viral injections

Virus AAV1-syn-GCaMP6s-WPRE-SV40 (400nl; 2E+13 vg/mL) and AVVdj-CaMKIIa-C1V1(E162T)-TS-P2A-mCherry-WPRE (200nl; titer 2.7e13 vg/mL) were injected simultaneously into layer 2/3 of left primary visual cortex (2.5 mm lateral and 0.3 mm anterior from the lambda, 200 µm from pia) using borosilicate pulled pipettes (tip diameter 2μm). 40-60% of the cells co-expressed both viruses. Virus mixture was injected at a rate of 80 nl/min, after all the volume was injected the pipette was hold for 5 minutes in the injection site to avoid flow back of the viruses due to pipette removal.

### Headplate procedure

3 weeks after virus injection mice were anesthetized with isoflurane (1-2%) and a custom designed titanium head plate was attached to the skull using dental cement in sterile conditions. Body temperature was maintained at 37 °C with an electric heater and monitored using a rectal probe. Dexamethasone sodium phosphate (2 mg/kg) and enrofloxacin (4.47 mg/kg) were administered subcutaneously. Carprofen (5 mg/kg) was administered intraperitoneally. A reinforced thinned skull window for chronic imaging (2 mm in diameter) was made above the injection site using a dental drill. A 3-mm circular glass coverslip was placed and sealed using a cyanoacrylate adhesive (Drew et al., 2010). During the surgery eyes were moisturized with eye ointment. After surgery animals received carprofen injections for 2 days as post-operative pain medication. Mice were allowed to recover for 5 days with food and water *ad libitum.*

### Behavioral system

We used a custom made treadmill attached to an angular position magnetic sensor. The water is delivered using a solenoid valve attached to a gravity water system. The waterspout was located at 1.5 mm from the animal’s mouth. The volume delivered for each correct trial was 4μl determined by the opening duration of the solenoid valve. Licking was monitoring with a commercial capacitive touch sensor attached to the waterspout. All signals were recorded to a host computer using a Digital Acquisition Board using MATLAB. An Arduino Uno connected via an USB interface to the host computer controlled visual stimulation and water delivery.

### Visual stimulation

Visual stimuli were generated using MATLAB Psychophysics Toolbox and displayed on a LCD monitor positioned 15 cm from the right eye at 45° to the long axis of the animal. Visual stimuli consisted of full-field sine wave drifting-gratings (contrasts: 100%, 50% and <40%, 0.035 cycles/°, 2 cycles/sec) drifting in two orthogonal directions presented for 2 sec, followed by 6 sec of mean luminescence. Experiments in the absence of visual stimuli (Fig. 8) were recorded with the monitor displaying a gray screen with mean luminescence similar to drifting-gratings.

### Behavioral training

After recovery from headplate implantation mice were weighted and handled for 2 days under water restriction until they reach 85% of their original weight, during this time mice underwent an habituation training to lick the waterspout and maneuver on the treadmill for 15-30 minutes daily. One hour before behavioral training food was removed. After the habituation period mice underwent a training phase for 3 days consisting in one session of 200 trials where water reward was automatically delivered following the Go signal (contrast 100%). Licking during the No-Go signal was punished with high frequency noise (200Hz). Following the training phase mice licked preferentially in water reward periods and avoided licking in No-Go periods. After the training phase the task phase began (day 1) where Go and No-Go visual stimuli (contrast 50%) were presented randomly in two sessions of 150 trials each separated by 10 min. Each stimulus was presented 50% of the time avoiding the presentation of the same stimulus more than two times in a row. After 7 days of the task phase mice reached a performance level above 75% that plateau for at least 8 days (Fig. 1C). Daily water supplementation was done to keep weight at 85% of the original value before animals were kept in their home cages overnight where food was available *ad libitum*.

Performance was calculated during the task phase as P= hits/(hits+miss) – false choices/(false choices+correct rejects).

### Simultaneous two-photon calcium imaging and photostimulation

Imaging experiments were preformed 7-28 days after head plate fixation. During recording sessions mouse is awake (head fixed) and can move freely on a treadmill. The imaging setup and the objective were completely enclosed with blackout fabric and a black electrical tape to avoid light contamination leaking into the PMTs. We used calcium imaging to monitor the activity of neuronal populations (Yuste and Katz, 1991). Two-photon imaging and optogenetic photostimulation were performed with two different femtosecond-pulsed lasers attached to a commercial microscope. An imaging laser (Ti:sapphire; λ = 940 nm) was used to excite a genetically encoded calcium indicator (GCaMP6s) while a photostimulation laser (low repetition rate pulse-amplified laser; λ = 1040 nm) was used to excite a red shifted opsin (C1V1) that preferentially responds to longer wavelengths (Packer et al., 2012). The power of both lasers was controlled by two independent pockels cells.

The two laser beams on the sample are individually controlled by two independent sets of galvanometric scanning mirrors. The imaged field of view was ∼240X240 µm (25X NA 1.05 XLPlan N objective), comprising 50-120 neurons. Neuronal contours were automatically identified using independent component analysis and image segmentation (Mukamel et al., 2009). Short movies (∼720 s) with a sample rate of 200-250 ms/frame were collected at time intervals of 5-10 min for up to 2h (Imaging laser power<50 mW; dwell time 2 μs/pixel; 256X256 pixels in the whole field of view).

Population photostimulation was performed splitting the laser beam into multiple foci using holographic stimulation through a Spatial Light Modulator (SLM). We adjusted the power of photostimulation in each neuron (Photostimulation laser power ∼5 mW) such that the amplitude of calcium transients evoked by C1V1 activation was not significantly different to the amplitude of calcium transients evoked by visual stimulation with drifting-gratings as previously shown (Carrillo-Reid et al., 2016). Single cell photostimulation was performed with a spiral pattern scanned by a pair of post-SLM galvanometric mirrors delivered from the center of the cell to the boundaries of the soma at 0.001 pix/μs (12μm diameter; 20 Hz) for one second. Photostimulation began 50ms after the onset of visual stimuli. The pulse repetition rate for photostimulation laser was 1MHz.

Simultaneous imaging and photostimulation was controlled by Prairie View and custom made software running in MATLAB.

For imaging experiments during behavioral task we performed 250 trials divided in 10 sessions (25 trials each) separated by 5 minutes. The first 3 sessions and the last 3 sessions were discarded from the analysis to avoid underestimation of behavioral performance due to motivation factors.

### Image processing

Image processing was performed with Image J (v.1.42q, National Institutes of Health) and custom made programs written in MATLAB as previously described (Carrillo-Reid et al., 2008; Carrillo-Reid et al., 2016; Cossart et al., 2003b). Acquired images were processed to correct motion artifacts using TurboReg. Regions of interest (ROIs) representing neurons were automatically identified using principal component analysis (PCA) and independent component analysis (ICA) algorithms written in Matlab (Mukamel et al., 2009). Calcium transients were computed as changes in fluorescence: (F_i_ – F_o_)/F_o_, where F_i_ denotes the fluorescence intensity at any frame and F_o_ denotes the basal fluorescence of each neuron (Miller et al., 2014). Spikes were inferred from the gradient (first time derivative) of calcium signals using a threshold of 3 standard deviations (S.D) above noise. We constructed an *N* × *F* binary matrix, where *N* denotes the number of active neurons and *F* represents the total number of frames for each movie. Each row in the binary matrix represents the activity of one neuron. To visualize neuronal activity the binary matrix was plotted as a raster plot where ones are represented by dots.

### Identification of neuronal ensembles

To identify neuronal ensembles from population calcium imaging recordings we constructed multidimensional population vectors that contain the information of the simultaneous activity of recorded neurons. The method is based on vectorial analysis (Carrillo-Reid et al., 2017; Carrillo-Reid et al., 2015a). Only population vectors with more active neurons in a given time than the ones expected by chance (P < 0.01) were considered for analysis. We tested the significance of population vectors against the null hypothesis that the synchronous firing of neuronal pools is given by a random process (Shmiel et al., 2006). Such population vectors can be used to compare the network activity as a function of time in different experimental conditions (Brown et al., 2005; Carrillo-Reid et al., 2008; Sasaki et al., 2007; Schreiber et al., 2003; Stopfer et al., 2003). The number of dimensions for each experiment is given by the total number of recorded cells. The temporal vectorization of the network activity allows the discrimination of specific coactive groups that are repeated at different times (Brown et al., 2005; Carrillo-Reid et al., 2008; Schreiber et al., 2003). To measure the similarity between population vectors at different experimental conditions we computed the normalized inner product (Carrillo-Reid et al., 2008; Sasaki et al., 2007; Schreiber et al., 2003), which represents the cosine of the angle between two vectors. To identify neuronal ensembles we constructed similarity maps from all the possible combinations of similarity values between vector pairs. In this way the time course of each neuronal ensemble is defined by each factor of the singular value decomposition (SVD) of the binary similarity map. The factorization is defined by a symmetric matrix M=V∑V^T^, where V and V^T^ are orthonormal and the elements of ∑ denote the singular values. The factors from the SVD associated with a singular value whose magnitude was above chance level represent the population vectors when a recurrent ensemble was active as previously published (Carrillo-Reid et al., 2017; Carrillo-Reid et al., 2015a; Carrillo-Reid et al., 2015b; Carrillo-Reid et al., 2016). To determine if the representative population vectors that define cortical ensembles could appear by chance we shuffled the overall activity matrix preserving the dimensionality of population vectors and compared the probability distribution of similarity coefficients from real data and shuffled data.

### Identification of neurons with pattern completion capability

To identify the neurons to be targeted by two-photon optogenetics we used conditional random fields (CRFs) to model the conditional probability distribution to see a given neuronal ensemble firing together (Carrillo-Reid et al., 2017). We used CRFs to capture the contribution of specific neurons to the overall network activity defined by population vectors belonging to a given neuronal ensemble. We generated a graphical model where each node represents a neuron in a given ensemble and edges represent the dependencies between neurons. 90% of the recorded data were used for training and the remaining 10% were used for cross-validation. The model parameters were determined by the local maximum of the likelihood function in the parameter space. Based on the model the node strength between adjacent nodes is defined by the summation of the edge potentials representing concomitant activity between neurons. The defined node strength reflects the conditional probability of co-activation between neurons. To measure which neurons are the most important for a given ensemble we computed the standard receiver operating characteristic curve (ROC), taking as ground truth the timing of a particular visual stimuli. The computation of the area under the curve (AUC) from the ROC curve that represents the performance of each neuron and the node strength that represents the connectivity between adjacent nodes were used to capture in a two dimensional space the most important neurons from each ensemble. As it has been shown recently, high ranked neurons observed in this two dimensional space have the potential to recall a given ensemble. CRF models were trained using the Columbia University Yeti Shared HPC cluster. The code used for CRF models can be found at https://github.com/hanshuting/graph_ensemble.

